# Comparing faster evolving *rplB* and *rpsC* versus SSU rRNA for improved microbial community resolution

**DOI:** 10.1101/435099

**Authors:** Jiarong Guo, James R. Cole, C. Titus Brown, James M. Tiedje

## Abstract

Many conserved protein-coding core genes are single copy and evolve faster, and thus are more resolving phylogenetic markers than the standard SSU rRNA gene but their use has been precluded by the lack of universal primers. Recent advances in gene targeted assembly methods for large shotgun metagenomes make their use feasible. To evaluate this approach, we compared the variation of two single copy ribosomal protein genes, *rplB* and *rpsC*, with the SSU rRNA gene for all completed bacterial genomes in NCBI RefSeq. As expected, among pairwise comparisons of all species that belong to the same genus, 94.9% and 91.0% of the pairs of *rplB* and *rpsC*, respectively, showed more variation than did their SSU rRNA gene sequences. We used a gene-targeted assembler, Xander, to assemble *rplB* and *rpsC* from shotgun metagenomic data from rhizosphere samples of three crops: corn (annual), and *Miscanthus* and switchgrass (both perennials). Both protein-coding genes separated all three communities whereas the SSU rRNA gene could only separate the annual from the two perennial communities in ordination analyses. Furthermore, assembled *rplB* and *rpsC* yielded significantly higher numbers of OTUs (alpha diversity) than the SSU rRNA gene. These results confirm these faster evolving marker genes offer increased resolution of for comparative microbiome studies.

## Introduction

Shaped by 3.5 billion years of evolution, microorganisms are estimated to comprise up to one trillion species and the majority of genetic diversity in the biosphere (Locey and Lennon 2016). Our understanding of this diversity is limited by the magnitude of extend species, and that the majority are yet to be cultured and their physiology or functions characterized. Since the pioneering work of Carl Woese in the late 1970s, the small subunit (SSU) rRNA gene has been the dominant marker used in microbial community structure analyses (Woese and Fox 1977; Lane *et al*. 1985; Huse *et al*. 2008; Caporaso *et al*. 2012). Although it has advanced our understanding of the microbial world, it does have important limitations, namely that it is highly conserved and that there are usually multiple copies, and some with intra-genomic variations, making this gene problematic for taxonomic identification at species and ecotypes levels and, in some cases, incapable of reflecting community distinctions at ecologically meaningful levels (Case *et al*. 2007; Wu and Eisen 2008; Roux *et al*. 2011).

With the accelerated accumulation of microbial genomes in recent years (Land *et al*. 2015), whole genome-based comparison is now feasible and a more accurate method for species and strain identification (Goris *et al*. 2007; Luo *et al*. 2011; Scortichini *et al*. 2013; Varghese *et al*. 2015; Rodriguez-R *et al*. 2018a, 2018b). However, whole genome-based comparison is relatively expensive computationally, compared to marker genes, and it is not yet possible to obtain genome sequences of many members of natural microbial communities due to the challenge of metagenome assembly from complex environmental communities (Howe *et al*. 2014; Li *et al*. 2015; Olson *et al*. 2017). Hence, marker gene analysis remains useful. Single copy protein coding housekeeping genes stand out as the best candidates as mark genes. First, their single copy status provides more accurate species and strain counting, identification and OTU (Operational Taxonomic Unit) clustering than the SSU rRNA gene. Second, they are present in virtually all known members of the three domains of life. Third, protein coding genes evolve faster than ribosomal RNA genes not only because rRNA genes are more conserved due to their more critical role in ribosome function (compared to ribosomal proteins) (Carter *et al*. 2000), but also because of the redundancy in the genetic code, especially at the third codon position (Case *et al*. 2007).

Here, we evaluate two single copy protein coding genes, *rplB* (encoding 50S ribosomal large subunit protein L2), and *rpsC* (encoding 30S ribosomal small subunit protein S3) as potential housekeeping genes for phylogenetic markers for microbial community analyses. Earlier studies showed the potential of protein coding genes over SSU rRNA genes as higher resolution phylogenetic markers for microbial diversity analyses using both genomic data (111 genomes) and metagenomic data (< 6 Gbp by Sanger sequencing) (Case *et al*. 2007; Roux *et al*. 2011). We revisited this comparison with the now much larger data set - all completed bacterial genomes (~4500 with one contig) and then tested the resolving power of these two genes versus the SSU rRNA gene among different crop rhizospheres using large shotgun metagenomic data (~1TB). The novelty of our analyses is 1) gene-targeted assembly can recover single copy protein coding genes more efficiently and accurately than generic assembly tools from shotgun metagenomic data (Wang *et al*. 2015) and 2) the use of *de novo* OTU-based diversity analyses, commonly used in microbial diversity analyses, rather than just taxonomic identification that is limited by reference databases (Case *et al*. 2007; Roux *et al*. 2011). Our analyses also have advantages over existing tools such as phylOTU (Sharpton *et al*. 2011) and singleM (https://github.com/wwood/singlem) that generate *de novo* OTUs from short reads since short reads may not overlap and thus clustering could be unreliable.

## Methods

Bacterial genome assembly information from NCBI (ftp://ftp.ncbi.nlm.nih.gov/genomes/refseq/bacteria/assembly_summary.txt) was used to construct the link to download each genome based on the instructions described in this link (http://www.ncbi.nlm.nih.gov/genome/doc/ftpfaq/#allcomplete). Command line “wget” was then used to retrieve the genome sequences with links obtained from the above step.

For extracting genes from genomes, the SSU rRNA gene HMM (Hidden Markov Model) from SSUsearch (Guo *et al*. 2015) was used to recover rRNA genes. Aligned *rplB* and *rpsC* nucleotide sequences of the “training set” retrieved from the RDP FunGene database (Fish *et al*. 2013) were used to build the HMM models using hmmbuild command in HMMER (version 3.1b2) (Eddy 2009). The nhmmer command in HMMER was then used to identify SSU rRNA, *rplB* and *rpsC* sequences from bacteria genomes obtained from NCBI using score cutoff (-T) of 60. Next, nhmmer hits of least 90% of the length of the HMM model were accepted as the target gene. For the purpose of comparing SSU rRNA, *rplB* and *rpsC* gene distances, one copy of the SSU rRNA gene was randomly picked from each genome. Pairwise comparison among gene sequences was done using vsearch (version 1.1.3) with “--allpairs_global --acceptall --iddef 2” (Rognes *et al*. 2016). Two species of environmental interest, *Rhizobium leguminosarum* and *Pseudomonas putida*, and their genera and orders, were chosen for closer comparison of *rplB, rpsC*, and SSU rRNA gene distances.

The shotgun data are from DNA from seven field replicates of rhizosphere samples of three biofuel crops: corn (C) *Zea maize*, switchgrass (S) P*anicum virgatum*, and *Miscanthus* × gigantus (M) that had been grown for five years. Shotgun sequence data (150 bp paired ends sequenced from 275bp libraries) for the 21 samples were downloaded from the JGI web portal (http://genome.jgi.doe.gov/); JGI Project IDs are listed in Table S1. Raw reads were quality trimmed using fastq-mcf in EA-Utils (verison 1.04.662) (http://code.google.com/p/ea-utils) “-l 50 -q 30 -w 4 -k 0 -x 0 --max-ns 0 -X”. Overlapping paired-end reads were merged by FLASH (version 1.2.7) (Magoc and Salzberg 2011) with “-m 10 -M 120 -x 0.20 -r 140 -f 250 -s 25” as described in (18).

SSU rRNA gene (*E.coli* position: 515 - 806, V4) amplicon data amplified from the same DNA as shotgun sequence (forward primer: GTGCCAGCMGCCGCGGTAA, reverse primer: GGACTACHVGGGTWTCTAAT) were also downloaded from JGI portal (JGI project ID: 1025756) and trimmed the same way as shotgun data (described above). Paired ends were joined by FLASH (-m 10 -M 150 -x 0.08 -p 33 -r 200 -f 300 -s 25) (Magoc and Salzberg 2011) and primer sequences were removed by cutadapt (-f fasta --discard-untrimmed) (Martin 2011). For community analyses, the QIIME (version 1.8.0) open reference OTU picking method with a default OTU clustering distance cutoff of 0.03 for de novo clustering step was used for clustering and core_diversity_analyses. py was used for generating the OTU table with taxonomy information (Kuczynski *et al*. 2012). SILVA reference database 108 was used for taxonomy assignment and alignment. Each sample was subsampled to 57418 reads and singletons were removed (details in https://github.com/jiarong/2016-rplB).

For SSU rRNA gene analyses with shotgun data, SSU rRNA gene fragments and those aligned to a part of V4 region (*E. coli* position: 577 - 727) of each sample were identified using the SSUsearch pipeline (version 0.8.2) (Guo *et al*. 2015) and clustered using RDP’s McClust tool (Cole *et al*. 2014) at a distance of 0.05 and minimal overlap of 25 bp and minimum read length of 100 bp, following the tutorial in SSUsearch (http://microbial-ecology-protocols.readthedocs.io/en/latest/SSUsearch/overview.html).

Both *rplB* and *rpsC* sequences were assembled using Xander with “MAX_JVM_HEAP=500G, FILTER_SIZE=40, K_SIZE=45, genes = *rplB* and *rpsC*, MIN_LENGTH=150, THREADS=9” (Wang *et al*. 2015). The minimal length cutoff is set to 150 amino acids (450 bp in nucleotide). Sequence reads for each crop were assembled separately. The assembled *rplB* or *rpsC* sequences (nucleotide and protein) from the three crops were pooled and clustered using RDP’s McClust tool (Cole *et al*. 2014). For each gene, a table of OTU counts of each sample was made based on mean k-mer coverage of the representative sequence of each OTU (provided in “*_coverage.txt” output file from Xander). An implementation of this pipeline is publicly available at https://doi.org/10.5281/zenodo.1438073.

Further, the above OTU tables were loaded into R, singletons were removed, and all samples were rarefied to the sample with the least total read number using “rrarefy” in vegan package (Oksanen *et al*. 2015). Diversity analyses were done with the vegan package using function “rda” for ordination with Bray-Curtis distance index, “diversity” for Shannon diversity, “estimateR” for Chao1 species richness, and “specnumber” for OTU number, from the OTU (count) tables (details in https://github.com/jiarong/2016-rplB). For alpha diversity comparisons among genes, all samples were subsampled to 1200 sequences.

To assess how many potential target gene reads of *rplB* and *rpsC* were assembled by Xander, we did a six-frame translation of the short reads (nucleotide sequences) into protein sequences by transeq in EMBOSS tool (Rice, Longden and Bleasby 2000). We then searched HMMs against the protein sequences and the hits with bit score > 40 (e-value < 6.2 * 10^−6^) were treated as reads from the target gene. In addition, “*_match_reads.fa”, a collection of reads that share a k-mer (k=45) with assembled sequences, output from Xander, provided the reads assembled by Xander. Then we compared the fold coverage of reads found by hmmsearch and reads used by Xander, by estimating fold coverage of each read with median kmer coverage using khmer package (Brown *et al*. 2012; Crusoe *et al*. 2015).

## Results

A total of 4,457 of complete bacteria genomes (genome as one sequence) were downloaded, and 4,440 of them have all three genes. SSU rRNA gene copy number ranged from 1 to 16 with a mean of 4, and *rplB* and *rpsC* are single copy in 99.9% of genomes (Table S2). When evaluating intra-genomic variation among copies of SSU rRNA genes in completed genomes of *R. leguminosarum*, *P. putida* and *E. coli*, *E. coli* had the largest variation with a minimum of 95.4% identity (Fig. S1).

For the selected taxa, Rhizobiales, Pseudomonadales, *Rhizobium*, and *Pseudomonas*, *rplB* and *rpsC* had similar variations and both had larger variation among the genomes than SSU rRNA genes within their corresponding order (among genera), and genus (among species) (Fig. 1 and 2). When comparing all species of completed genomes that belong to the same genus, we found SSU rRNA gene has an identity range of 63.2% to 100.0% and a median of 95.2%, *rplB* has an identity range of 43.2% to 100.0% and a median of 87.2%, *rpsC* has an identity range of 46.0% to 100.0% and a median of 90.3%. Between *rplB* and SSU rRNA gene, 88,993 pairs (94.9% of total) had larger variation in *rplB*, 3,573 pairs (3.8%) had larger variation in SSU rRNA gene, and 1,167 pairs (1.2%) had the same variation (Fig. 3A); Between *rpsC* and SSU rRNA gene, 77,885 pairs (91.0% of total) had larger variation in *rpsC*, 6,074 pairs (7.1%) had larger variation in SSU rRNA gene, and 1,622 pairs (1.9%) had the same variation (Fig. 3B); Between *rplB* and *rpsC*, 54,755 pairs (63.7%) had larger variation in *rplB*, 28,393 pairs (33.0%) had larger variation in *rpsC* gene, and 2,808 pairs (3.3%) had the same variation for *rplB* and *rpsC* (Fig. 3C).

**Figure 1:**
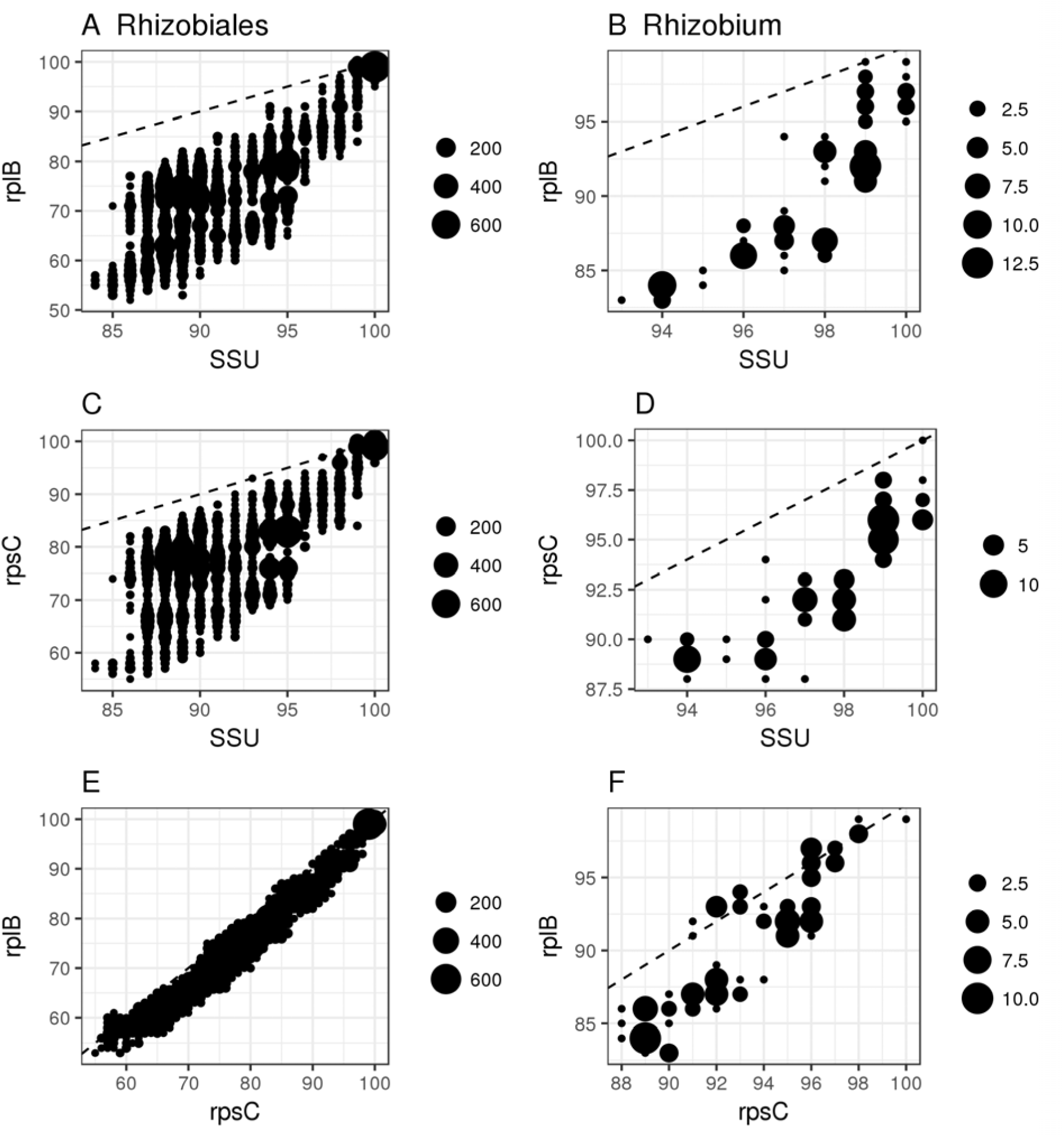
Pairwise comparisons among all genomes of Rhizobiales (panels A, C, E) and of all *Rhizobium* (panels B, D, F) using the SSU rRNA gene, *rplB* and *rpsC*. SSU rRNA gene identities are higher than *rplB* and *rpsC*, and *rplB* and *rpsC* have similar sequence identities in most genome pairs. The dashed line is y = x. Data below y=x line indicate the gene on X axis is more conserved. The dot size indicates the number of pairwise comparisons with those values.

**Figure 2:**
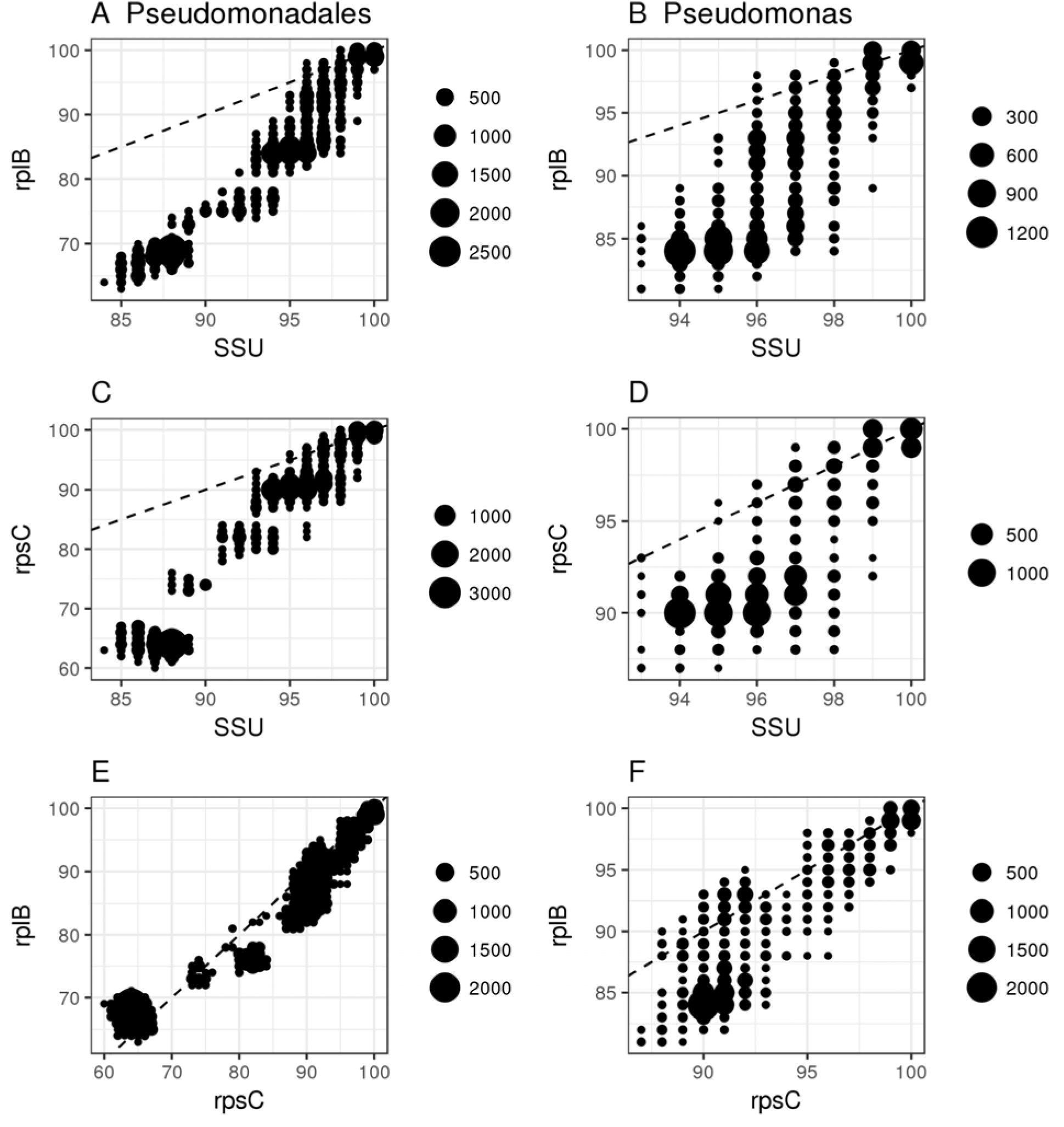
Pairwise comparison among all genomes of Pseudomonadales (panels A, C, E) and of all *Pseudomonas* (panels B, D, F) using the SSU rRNA gene, *rplB* and *rpsC*. SSU rRNA gene identities are higher than *rplB* and *rpsC* in most genome pairs, and *rplB* and *rpsC* have similar sequence identities. The dashed line is y = x. Data below y=x line indicate gene on X axis is more conserved. The dot size indicates the number of pairwise comparisons with those values.

**Figure 3:**
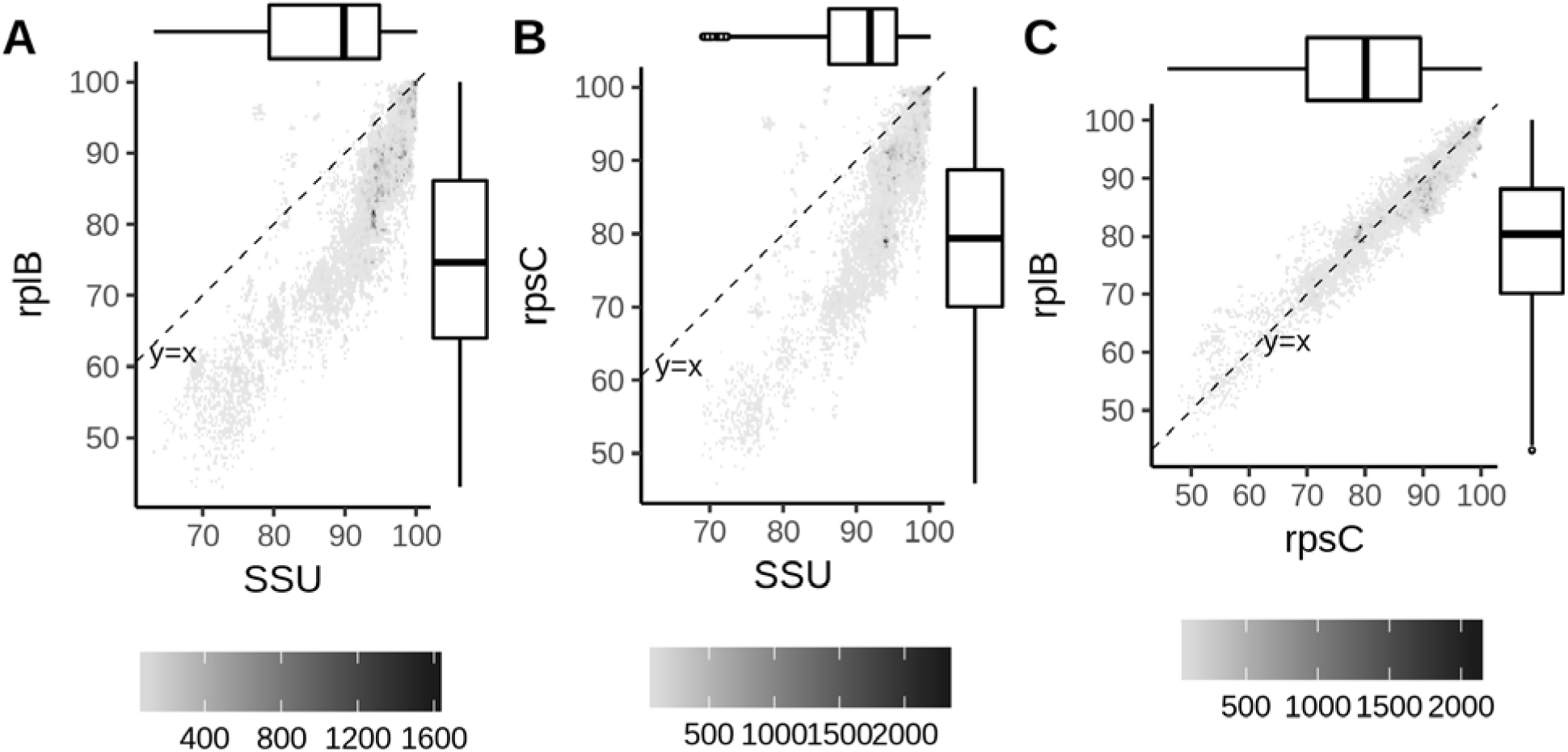
Pairwise comparison among all completed genomes of species in the same genus using the SSU rRNA gene, *rplB*, and *rpsC*. SSU rRNA gene identities are larger than *rplB* in most genomes. The diagonal dashed line is y = x and data below the line indicates the gene on X axis is more conserved. The dot intensity is the number of comparisons with those values. Subplot A, B, and C are comparisons of *rplB* and SSU rRNA gene (93,733 pairwise comparisons), *rpsC* and SSU rRNA gene (85,581 pairwise comparisons), *rplB* and *rpsC* (85,956 pairwise comparisons).

We compared SSU rRNA genes (V4) with *rplB and rpsC* to test the ability of shotgun data to resolve community differences among plant rhizospheres. We chose these two genes as they had a suitable length for Xander assembly (~ 660bp for *rpsC* and 830bp for *rplB*), were long enough for resolving power, and had HMMs that were both specific and sensitive for fragment recovery due to their uniqueness in sequence as parts of the ribosome. Both have also been used as phylogenetic markers in other shotgun metagenomic studies (Hug *et al*. 2013; Sharon *et al*. 2015). On average, 0.04% of total reads were identified as SSU rRNA gene fragments and 0.004% of total reads aligned to the 150 bp of V4 region of the gene with SSUsearch (Guo *et al*. 2015). Another 0.01% and 0.008% of total reads were identified as *rplB* and *rpsC*, respectively, by Xander (Table S3). The assembled sequences (using a length cutoff of 450 bp) have lengths ranging from 453 to 843bp with a median of 822bp for *rplB* and from 453 to 678bp with a median of 648 bp for *rpsC*. To test the sensitivity of Xander, we found that the number of potential *rplB* and *rpsC* reads assembled were 49.5% and 47.9%, respectively, of those defined by hmmsearch with bit score cutoff of 40 (Table S4) and have much higher fold coverage than the rest of reads in hmmsearch hits (excluding shared reads with Xander) (Figure S2).

Beta diversity analyses of all three genes showed that the rhizosphere communities of the annual crop, corn, were different from those of the two perennial grasses, *Miscanthus* and switchgrass, but only *rplB* and *rpsC* distinguished the communities of the two perennial grasses (Fig. 4). This was true whether the analysis was at the nucleotide or protein level. The alpha diversity of the corn rhizosphere communities was significantly lower than those of *Miscanthus* and switchgrass rhizospheres by all three measures except for Chao1 index with *rpsC* and SSU rRNA gene (Fig. 5). When comparing among genes, the numbers of OTUs from *rplB* and *rpsC* were also significantly higher than from the SSU rRNA gene (Fig. 5). Since SSUSearch returns shorter fragments (~100bp to 150bp) than Xander assembled genes (~ 450bp to 849bp), we also evaluated whether the longer fragments of SSU rRNA from amplicon data (~ 250 to 300 bp), could distinguish the two perennial grass communities, and they could not (Fig. 4).

**Figure 4:**
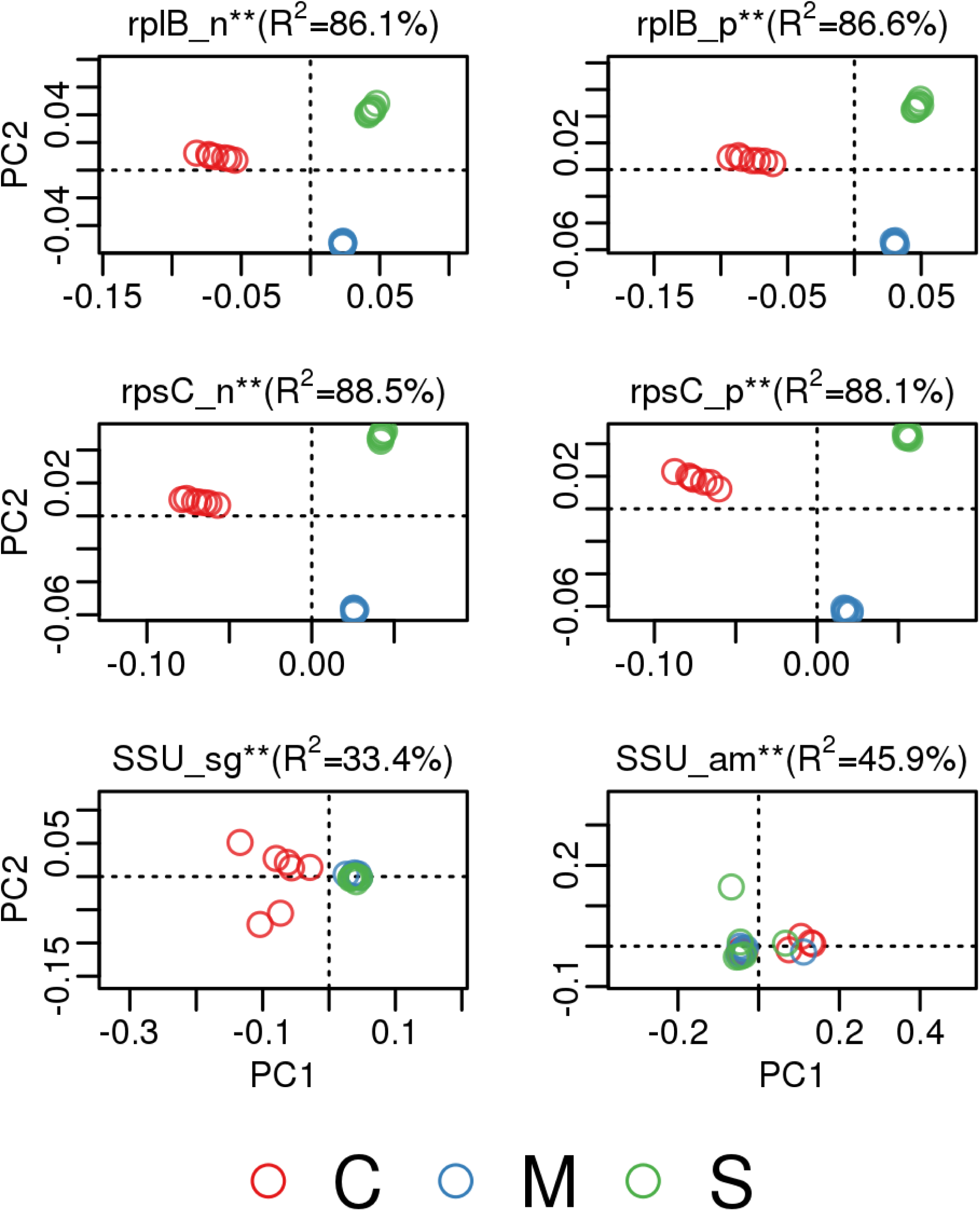
Comparison of the SSU rRNA gene, *rpsC* and *rplB* in beta diversity analyses (ordination) using large soil metagenome sequences from seven field replicates. All genes show that the microbial community of the corn (C) rhizosphere is significantly different from those of *Miscanthus* (M) and switchgrass (S) while *rplB* and *rpsC* at both the nucleotide (n) and protein (p) levels separate microbial communities of *Miscanthus* and switchgrass. The SSU rRNA gene does not separate *Miscanthus* and switchgrass with either shotgun (SSU.sg) or amplicon (SSU.am) data. “**” indicates p < 0.01 in PERMANOVA test.

**Figure 5:**
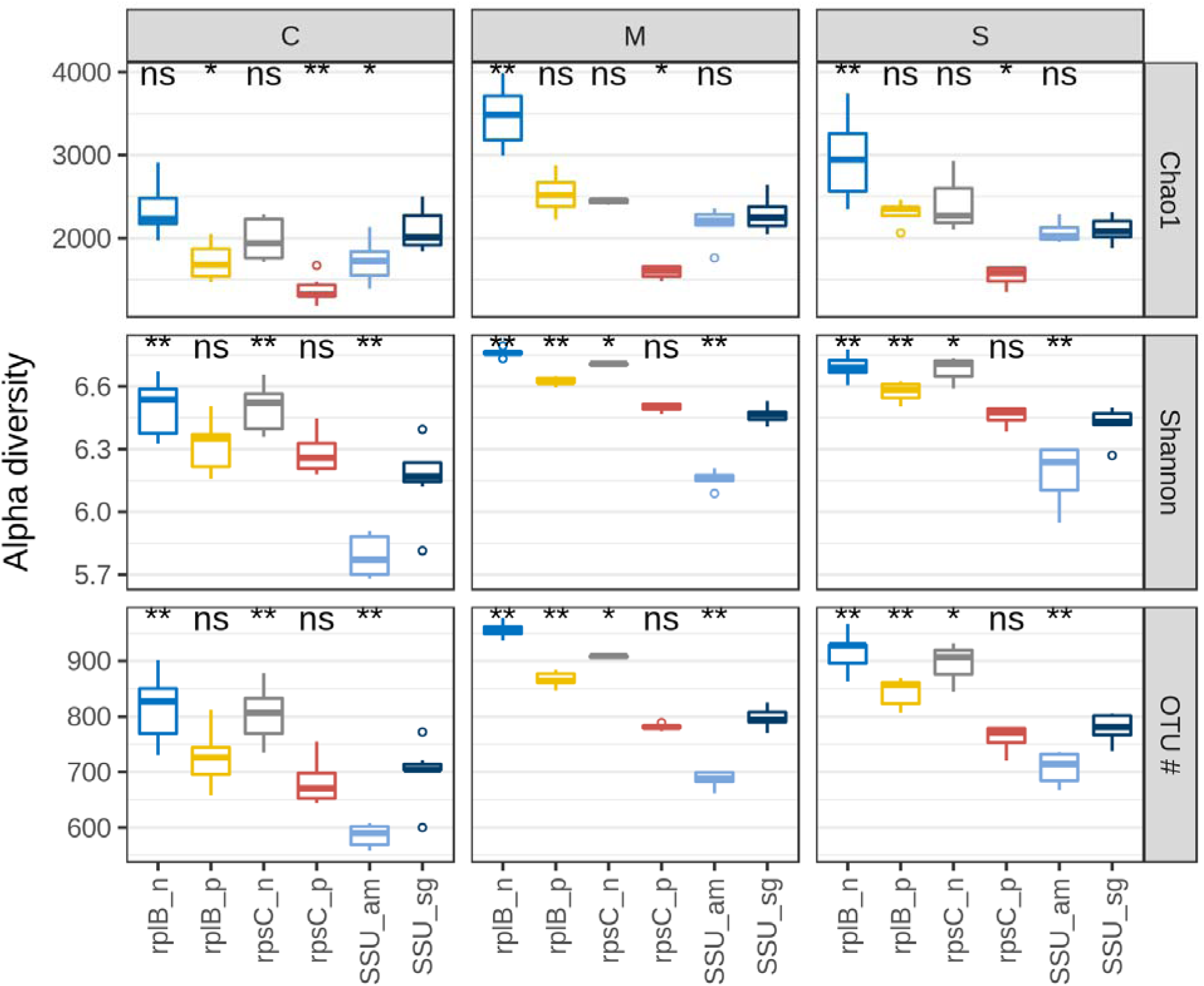
Comparison of the SSU rRNA gene, *rplB*, and *rpsC* in alpha diversity analyses (Chao1, Shannon, and OTU number) using large soil metagenome sequence. The labels with suffixes “_n” and “_p” means assembled nucleotide and protein sequences by Xander respectively; those with suffixes “_am” and “_sg” are SSU rRNA gene from amplicon data and shogun data respectively. All genes show that the microbial community of the corn (C) rhizosphere has significantly less alpha diversity than those of *Miscanthus* (M) and switchgrass (S) except for *rpsC* and SSU rRNA gene with Chao1 index (p < 0.01). Wilcoxon test was used to compare SSU_sg against each of the other genes including rplB_n, rplB_p, rpsC_n, rpsC_p and SSU_am. For Chao1 index, rplB_n and rpsC_n show significantly higher abundance than SSU_sg in *Miscanthus* and switchgrass; For Shannon index and OTU number, rplB_n and rpsC_n show significantly higher abundance than SSU_sg in all three crops (“***” is p < 0.001, ‘**’ is p < 0.01, ‘*’ is p < 0.05, “ns” is p > 0.05).

## Discussion

We confirmed the advantages of *rplB* and *rpsC* over the SSU rRNA gene as a more resolving phylogenetic marker using updated large genomic data (~4500 complete genomes) (Fig. 1, 2 and 3). We also demonstrated that *rplB* and *rpsC* can be assembled from large shotgun metagenomes and showed that they provided higher community resolution by separating *Miscanthus* and switchgrass rhizosphere samples while the SSU rRNA gene did not (Fig. 4). The two perennial grasses would be expected to have more similar microbiome than the annual since the latter is re-established each year while the fibrous perennial grass roots are more similar and not physically disturbed annually and thus do not have full regrowth at a new random site each year. In large genomic data analyses, *rplB* and *rpsC* show advantages in following three aspects:

First, SSU rRNA gene, a multiple copy gene, poses difficulties for interpreting species abundance, while *rplB* and *rpsC* do not have the same issue as it is single copy genes in > 99.9% of complete genomes (Table S2). Additionally, variations among multiple SSU copies can cause multiple OTUs (sequence clusters) from the same species (Figure S1) and thus leads to overestimation of species richness (31). Since a single copy of the *rplB* and *rpsC* genes is contained in every cell in a community, the abundance of *rplB* and *rpsC* gene sequences provides a reference for estimating the fraction of organisms possessing other genes and also average genome size (Raes *et al*. 2007; Frank and Sørensen 2011; Nayfach and Pollard 2015; Vital, Karch and Pieper 2017).

Second, *rplB* and *rpsC* are better able to differentiate closely related species based on their lower sequence identities compared to the SSU rRNA gene in pairwise comparisons among genomes (Fig. 1, 2, and 3). This is consistent with the crucial role SSU rRNA plays in translation (ensuring translation accuracy) (Carter *et al*. 2000), also confirmed by another study showing SSU rRNA genes (along with LSU rRNA genes, tRNA and ABC transporter genes) to be the most conserved genes (Isenbarger *et al*. 2008).

Third, SSU rRNA genes in genomes are also more prone to assembly errors (chimera) than single copy genes due to their higher overall nucleotide identity and the presence of highly conserved regions interspersed in SSU rRNA genes (Miller 2013; Sharon *et al*. 2015; Yuan *et al*. 2015). Note that these erroneous sequences might be further collected by databases and used as references for taxonomy, alignment, and chimera detection, and thus have an impact on common microbial ecology analyses. Single copy protein coding genes, however, are less challenging to assemble compared to the SSU rRNA gene since they have more unique sequences among different taxa and no highly conserved regions. They do however, require enough sequence for a sufficient number of gene assemblies (Hug *et al*. 2013; Sharon *et al*. 2015).

Additionally, this method provides for higher resolution community diversity analyses in large shotgun metagenomes, leveraging a scalable gene targeted assembler, Xander. Assembly is desirable for short read data to correctly identify the gene and provide enough length for resolving power, a major objective in ecology studies. While this method avoids the primer bias of amplicon sequencing (Farris and Olson 2007; Bergmann *et al*. 2011; Guo *et al*. 2015), it does miss the rarer species because not enough fragments of the targeted genes from rarer members are sequenced to allow their assembly, perhaps missing up to 50% of the cells of lowest abundance (Table S4 and Fig. S2). The hmmsearch though could also have recovered some false positives due to mistaken short-read identification and thus overestimated the total gene number (Orellana, Rodriguez-R and Konstantinidis 2017). However, *rplB* and *rpsC* still yielded significantly higher alpha diversity (Fig. 5) than the SSU rRNA gene despite missing rare members. These protein-coding genes, although shorter than the full-length SSU rRNA gene, are both more rapidly evolving and longer than produced with commonly used SSU rRNA gene primers. Thus they reveal more diversity among abundant members than SSU rRNA gene, which offsets and exceeds the diversity of the rare members that are not assembled, further confirming their higher resolution. Additionally, rare members are commonly removed in amplicon-based analyses, especially singletons, to remove spurious OTUs caused by sequencing error and to improve computational efficiency. Rare valid members are unavoidably removed in this process too, but this practice has been shown to have minimal impact on beta diversity analyses (Edgar 2013; Bokulich *et al*. 2013; Auer *et al*. 2017).

OTU distance cutoff could also impact community analyses in our comparison. We used a distance cutoff of 0.05 for *rplB* and *rpsC* since that is the cutoff recommend (22) (and we used) for SSUSearch, We used the standard, and more resolving 0.03 cutoff for the SSU rRNA gene amplicon data, but it showed less community resolution than *rplB* and *rpsC* indicating that our cutoff is appropriate and fairly evaluating the comparison (S1 text).

We did not attempt assembling the SSU rRNA gene because: i) as noted above, the conserved regions results in a high proportion of miss-assembly with short reads, generating chimeras (Miller 2013; Sharon *et al*. 2015; Yuan *et al*. 2015), ii) the existing tools, e.g. EMIRGE and REAGO, have drawbacks. EMIRGE relies heavily on reference databases and does not perform well when close relatives are absent. Further, EMIRGE was shown to have significant false positives (Fan, McElroy and Thomas 2012; Miller 2013; Yuan *et al*. 2015). REAGO has only been tested with low diversity, mock community data and does not report abundance information (Yuan *et al*. 2015).

We chose two protein-coding genes to ensure that our results were not gene specific, and both gave very similar results at both the nucleotide and protein levels. At least from extensive completed genomes, more than 99% of these two genes are single copy, making quantitative (ratio) comparisons with other genes consistent. For future use, *rplB* might have a slight advantage over *rpsC* since it is longer, about 830 bp on average vs. 660 bp of *rpsC* (Fish *et al*. 2013), providing a bit more resolving power, which is consistent with results in genome comparisons showing *rplB* has lower median sequence identity than *rpsC* (Fig 3).

Besides higher resolution in de novo OTU related analyses, it is of course possible to do taxonomy related analyses by finding the best match to the assembled sequence of these marker genes in reference databases and potentially finer taxonomic resolution than provided by SSU rRNA. But, the reference databases of *rplB* and *rpsC* are mostly from sequenced genomes and hence are very unbalanced and incomplete compared to SSU rRNA gene databases (Quast *et al*. 2013; Cole *et al*. 2014; Wang *et al*. 2015) so this use is not generally beneficial at this time.

Although the sequencing depth needed varies depending on community diversity, we provide an estimate based on our rhizosphere soil samples as a guide. The reads from *rplB* are around 0.01% of total (Table S3). Assuming a fold coverage of 3000 of *rplB* for each sample, to be comparable to 3000 amplicons in planning amplicon-based studies, one needs about 25 Gbp (3000 * 830 / 0.01%) of shotgun metagenome (830 bp is the average gene length of *rplB*). The major requirement for using this method beyond sufficient shotgun sequence depth is an access to a high performance computer since large memory (> 250 Gb recommended for soil samples) is needed for the deBruin graph (the most memory and cpu consuming step) to run Xander. However, once one has the deBruin graph, it can also be used to assemble other genes of interest (Wang *et al*. 2015).

## Conclusion

We demonstrated that *rplB* and *rpsC*, single copy protein coding genes can provide finer resolution of taxa and hence better distinguish among communities than the more commonly used SSU rRNA gene and also provide finer scale *de novo* OTU diversity analysis. This method does require shotgun sequence of sufficient depth, so is currently more costly than amplicon based analyses, but as sequencing costs decline, capacity and access increase, and read length and genome reference databases grow, single copy protein coding genes such *rplB* and *rpsC* have the potential to complement or even replace the SSU rRNA gene as a phylogenetic marker and better reflect the ecology of microbial communities.

## Author contributions

J.G. performed the analyses under the supervision of J.M.T., C.T.B. and J.R.C.. All also helped with the analysis approaches and writing of the manuscript.

## Acknowledgement

We thank Ribosomal Database Project (RDP), Institute for Cyber-Enabled Research (iCER) and High Performance Computing Center (HPCC) at Michigan State University for technical support. Support for this research was provided by the U.S. Department of Energy, Office of Science, Program of Biological and Environmental Research (Awards DE-FC02-07ER64494 and DE-FG02-99ER62848), and by the National Science Foundation DBI-1356380, DBI-1759892, and Long-term Ecological Research Program (DEB 1637653) at the Kellogg Biological Station, and by Michigan State University AgBioResearch.

