## supplemental materials for "Comparing faster evolving *rplB* and *rpsC* versus SSU rRNA for improved microbial community resolution"

### Supplemental Material

Table S1: Rhizosphere soil shotgun metagenomic data of seven replicates (1 to 7) for each crop (C = corn, M = *Miscanthus*, and S = switchgrass for all tables).

| Sample | JGI project ID | Data size (Gb) |
| --- | --- | --- |
| C1 | 1023764 | 46 |
| C2 | 1023767 | 39 |
| C3 | 1023770 | 57 |
| C4 | 1023773 | 53 |
| C5 | 1023776 | 57 |
| C6 | 1023779 | 51 |
| C7 | 1023782 | 46 |
| M1 | 1023785 | 56 |
| M2 | 1023788 | 57 |
| M3 | 1023791 | 50 |
| M4 | 1023794 | 42 |
| M5 | 1023797 | 45 |
| M6 | 1023800 | 43 |
| M7 | 1018623, 1018611 | 32 |
| S1 | 1023803 | 40 |
| S2 | 1018626, 1018614 | 28 |
| S3 | 1023806 | 44 |
| S4 | 1023809 | 49 |
| S5 | 1018629, 1018617 | 27 |
| S6 | 1023812 | 50 |
| S7 | 1023815 | 39 |

Table S2: Summary of the SSU rRNA gene, *rplB* and *rpsC* copy number in genomes.

| Copy | SSU | *rplB* | *rpsC* |
| --- | --- | --- | --- |
| 1 | 635 | 4443 | 4013 |
| 2 | 711 | 4 | 4 |
| 3 | 560 | 0 | 0 |
| 4 | 585 | 0 | 0 |
| 5 | 372 | 0 | 0 |
| 6 | 484 | 0 | 0 |
| 7 | 556 | 0 | 0 |
| 8 | 194 | 0 | 0 |
| 9 | 92 | 0 | 0 |
| 10 | 107 | 0 | 0 |
| 11 | 58 | 0 | 0 |
| 12 | 27 | 0 | 0 |
| 13 | 26 | 0 | 0 |
| 14 | 38 | 0 | 0 |
| 15 | 4 | 0 | 0 |
| 16 | 1 | 0 | 0 |
| 18 | 0 | 0 | 0 |
| 20 | 0 | 0 | 0 |

Table S3: Summary of reads and the proportion identified as the SSU rRNA gene, *rplB* and *rpsC* in soil metagenomes of seven replicates of each crop.

|  | Total Reads | SSU % | V4 % | *rplB* % | *rpsC* % |
| --- | --- | --- | --- | --- | --- |
| C1 | 232234095 | 0.050 | 0.005 | 0.013 | 0.010 |
| C2 | 220844998 | 0.043 | 0.004 | 0.010 | 0.007 |
| C3 | 282123335 | 0.042 | 0.004 | 0.012 | 0.009 |
| C4 | 260542662 | 0.041 | 0.004 | 0.011 | 0.008 |
| C5 | 285873232 | 0.050 | 0.005 | 0.012 | 0.009 |
| C6 | 250477617 | 0.050 | 0.005 | 0.012 | 0.009 |
| C7 | 262943930 | 0.043 | 0.004 | 0.011 | 0.008 |
| M1 | 274060925 | 0.037 | 0.004 | 0.010 | 0.007 |
| M2 | 278278868 | 0.036 | 0.004 | 0.010 | 0.008 |
| M3 | 244772969 | 0.038 | 0.004 | 0.010 | 0.008 |
| M4 | 206129129 | 0.034 | 0.003 | 0.010 | 0.007 |
| M5 | 225964704 | 0.033 | 0.003 | 0.010 | 0.007 |
| M6 | 215320045 | 0.034 | 0.003 | 0.010 | 0.008 |
| M7 | 160726636 | 0.044 | 0.004 | 0.009 | 0.007 |
| S1 | 192776425 | 0.035 | 0.003 | 0.010 | 0.008 |
| S2 | 143660127 | 0.040 | 0.004 | 0.009 | 0.007 |
| S3 | 218743879 | 0.035 | 0.003 | 0.010 | 0.007 |
| S4 | 249773480 | 0.031 | 0.003 | 0.009 | 0.007 |
| S5 | 140239637 | 0.036 | 0.003 | 0.009 | 0.006 |
| S6 | 254749228 | 0.032 | 0.003 | 0.010 | 0.007 |
| S7 | 194863138 | 0.035 | 0.003 | 0.010 | 0.007 |

Table S4. Evaluation of potential *rplB* and *rspC* gene reads used (assembled) by Xander. Fold coverage is calculated as Hits * 120/Gene length. The test dataset used here is C1 (198979842 reads in total and all reads trimmed to 120 bp). “Gene length” is the average of full gene lengths in bp.

|  | Method | Total | Hits | % | Gene length | Fold coverage |
| --- | --- | --- | --- | --- | --- | --- |
| *rplB* | xander | 198979842 | 14359 | 0.0072 | 830 | 2076 |
| *rpsC* | xander | 198979842 | 11548 | 0.0058 | 660 | 2100 |
| *rplB* | hmmsearch | 198979842 | 28697 | 0.0144 | 830 | 4149 |
| *rpsC* | hmmsearch | 198979842 | 24113 | 0.0121 | 660 | 4384 |

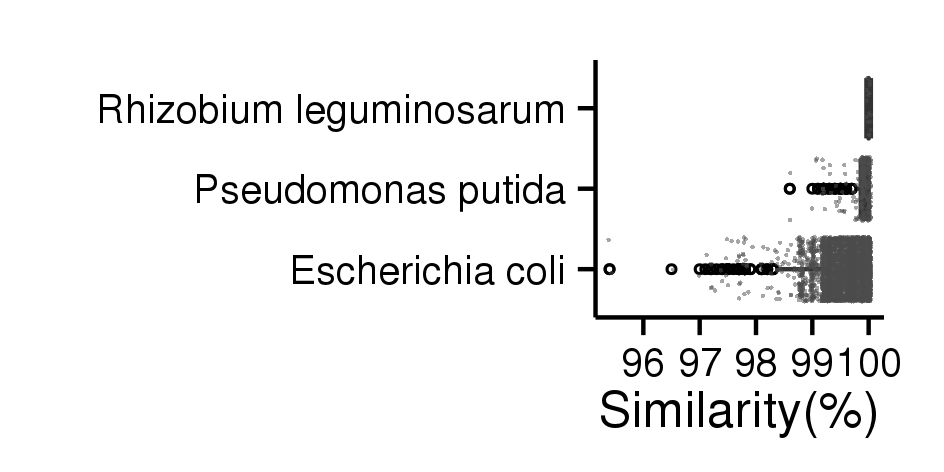

Figure S1: Intra-genome variation of SSU rRNA gene of three specie. Each dot is a pairwise comparison of SSU rRNA genes within the same genome. *R. leguminosarum* has 5 genomes with 15 comparisons*, P, putida* has 13 genomes with 227 comparisons, and *E. coli* has 150 genomes with 3201 comparisons. *E. coli* has largest intra-genome SSU rRNA gene variation among the three.

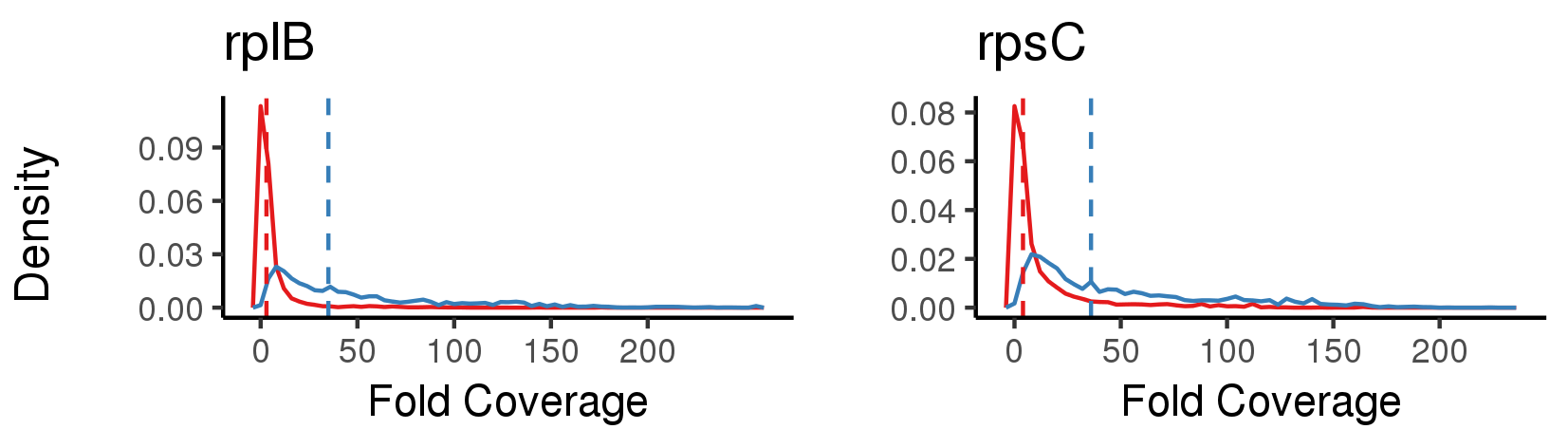

Figure S2. Comparison fold coverage of reads used in assemblies and read hits by hmmsearch (excluding those used in assemblies) for *rplB* and *rpsC*. X axis is fold coverage of reads. Y axis is the density (relative abundance of reads with a specific fold coverage) of the distribution of fold coverage of all reads. Reads used in assemblies (blue) have a much higher fold coverage than hits by hmmsearch (red). The dashed vertical lines are the median of all fold coverages.

S1 Text. Further evaluation of the impact of OTU clustering cutoffs on the method comparisons.

In our comparison of three genes’ resolution to separate the rhizosphere of three biofuel crops, a distance cutoff of 0.05 was chosen for the following reasons. Sequencing error has large impact on clustering with SSU rRNA gene fragments from shotgun data since the pairwise distance is calculated by dividing the number of mismatch by overlap positions in alignment with a minimum of 50bp in this case, and thus sequencing error could cause larger pairwise distance (e.g. one sequencing error over 50bp overlap could cause 0.02 distance) and more spurious clusters, and a distance cutoff of 0.05 is recommended by SSUsearch (Guo *et al.* 2015). Accordingly, 0.05 is used for *rplB* and *rpsC* for fair comparison. Further, 0.03 is used for SSU rRNA gene amplicon data since the average amplicon (Miseq) length is about 250 bp, so more sequencing error tolerant in terms of OTU clustering, i.e. one sequencing error only cause 0.004 distance. However, the results that amplicon SSU rRNA gene data have less resolution than shotgun SSU rRNA gene and *rplB* still stand since there would be less OTUs and thus less resolution if a distance cutoff of 0.05 was used in amplicon SSU rRNA gene data. We are also aware that amplicon sequence variant (ASV) based methods can provide higher resolution than OTU clustering based methods for amplicon SSU rRNA gene data. We choose OTU clustering-based method since ASV-based methods are only suitable for amplicon data and not for shotgun data (Callahan *et al.* 2016; Callahan, McMurdie and Holmes 2017).
